# Polarity sorting drives remodeling of actin-myosin networks

**DOI:** 10.1101/314484

**Authors:** Viktoria Wollrab, Julio M. Belmonte, Maria Leptin, François Nédeléc, Gijsje H. Koenderink

**Affiliations:** AMOLF, Science Park 104, 1098 XG Amsterdam; EMBL, Cell Biology and Developmental Biology Unit and Director’s Research Unit, Meyerhofstraße 1, Heidelberg

## Abstract

Cytoskeletal networks of actin filaments and myosin motors drive many dynamic cell processes. A key characteristic of these networks is their contractility. Despite intense experimental and theoretical efforts, it is not clear what mechanism favors network contraction over expansion. Recent work points to a dominant role for the nonlinear mechanical response of actin filaments, which can withstand stretching but buckle upon compression. Here we present an alternative mechanism. We study how interactions between actin and myosin-2 at the single filament level translate into contraction at the network scale by performing time-lapse imaging on reconstituted quasi-2D-networks mimicking the cell cortex. We observe myosin end-dwelling after it runs processively along actin filaments. This leads to transport and clustering of actin filament ends and the formation of transiently stable bipolar structures. Further we show that myosin-driven polarity sorting leads to polar actin aster formation, which act as contractile nodes that drive contraction in crosslinked networks. Computer simulations comparing the roles of the end-dwelling mechanism and a buckling-dependent mechanism show that the relative contribution of end-dwelling contraction increases as the network mesh-size decreases.

## Introduction

Cells have the remarkable ability to actively deform themselves to drive vital processes such as cell division, cell migration and multicellular tissue dynamics. The main determinant of cell shape is the actin cytoskeleton, which actively deforms the plasma membrane by generating pushing and pulling forces (Blanchoin et al., 2014). This activity relies on the structural polarity of actin filaments, which have structurally distinct ends denoted as the *minus* and the *plus end* (also referred to as the pointed and the barbed end). Hydrolysis of adenosine triphosphate (ATP) bound to actin monomers that add onto the plus end of growing filaments provides chemical energy, which allows actin filaments to exert polymerization forces at their plus ends. Myosin-2 motors take advantage of the structural polarity of F-actin to move in a directional manner toward the plus end, again using energy released from ATP hydrolysis. The motors work together in teams known as bipolar filaments, with motor heads on the two ends and the tails packed in the center. This bipolarity allows myosin filaments to slide anti-parallel actin filaments in opposing directions.

Together with accessory crosslinking proteins, actin and myosin-2 form different contractile assemblies in the cell (Murrell et al., 2015). Right underneath the plasma membrane, they form a thin and dense polymer mesh known as the cortex. A key function of the cortex is its contractility, which can drive global cell rounding as cells enter mitosis, local membrane constriction during cell division, and global tissue deformation in developing embryos (Salbreux et al., 2012). In the cytoplasm, actin and myosin form bundled contractile structures known as stress fibers in adherent cells and 3D-meshworks in large oocytes and early embryo cells (Field and Lénárt, 2011; Naumanen et al., 2008). Finally, during cytokinesis, actin and myosin form a contractile structure known as the cytokinetic ring, which constricts the membrane (Wollrab et al., 2016).

Despite detailed knowledge of the molecular composition of the actin-myosin ‘contractome’ (Biro et al., 2013; Zaidel-Bar et al., 2015), the molecular mechanism for the contractile activity of the actin-myosin cytoskeleton is unclear since the actomyosin bundles and meshworks in nonmuscle cells are disordered in terms of the filament orientations and polarities. This is completely unlike muscle sarcomeres, where the actin and myosin are arranged in repeating arrays with myosin bipolar filaments localized in-between antiparallel actin filaments having their minus ends inwards and their plus ends outwards. In this case, the localization of the myosin clusters in the vicinity of F-actin minus ends and the anchoring of the actin filament minus ends at the Z-discs convert the sliding activity of the motors into pure contraction (Gautel and Djinovic-Carugo, 2016). By contrast, motor-mediated sliding of rigid filaments in random networks is in principle equally likely to result in contraction or expansion (Belmonte et al., 2017; Mendes Pinto et al., 2012). Experimentally, both contraction and expansion have indeed been demonstrated in reconstituted networks of microtubules and motors (Foster et al., 2015; Sanchez et al., 2012; Torisawa et al., 2016) and in cells (Lu et al., 2013). In contrast, actomyosin assemblies in cells always contract, and reconstituted networks of actin filaments and myosin motors are also nearly always contractile (Stam et al., 2017).

It has been a long-standing question why actin-myosin networks are biased towards contraction and different microscopic mechanisms have been proposed (Alvarado et al., 2017). The currently most favored mechanism is based on the nonlinear elastic properties of actin filaments. Actin filaments are semiflexible polymers with a persistence length of around 10 μm (Kang et al., 2012). As a consequence, they resist stretching but readily buckle upon compression forces induced by molecular motors. Theoretical models of active networks predict that buckling will cause contraction in both bundles and meshworks, independent of filament polarity (Lenz et al., 2012; Ronceray et al., 2016). Effectively, myosin bipolar filaments interacting with crosslinked actin networks act as contractile force dipoles (Mackintosh and Levine, 2008). This buckling scenario is supported by direct experimental observations of filament buckling and telescopic contraction in reconstituted actin-myosin bundles and random networks (Linsmeier et al., 2016; Murrell and Gardel, 2012).

Yet, analytical and computational models predict that preferential contraction can also occur in the absence of filament buckling. The basic idea is that if motors processively walk along cytoskeletal filaments towards one end (the plus end in case of myosin-2) and dwell before they detach, they will cause polarity sorting of the filaments and eventually contraction of both bundles and networks (Kruse and Jülicher, 2000; Zumdieck et al., 2007). In networks, a clear signature of polarity sorting is the formation of radial arrays of filaments known as asters, where the filament ends point inwards and motors accumulate in the center. In case of microtubule systems, several motors have indeed been shown to exhibit end-dwelling behavior (Akhmanova and Hoogenraad, 2005) and cause the formation of polar asters (Foster et al., 2015; Surrey et al., 2001; Torisawa et al., 2016). While this phenomenon is now well accepted for microtubules (Tan et al., 2018), it is not known whether it can also operate for actin. In case of myosin-2, there are observations of processive motion along actin filaments in motility assays and in dense actin-myosin networks (Sellers and Kachar, 1990; Soares e Silva et al., 2011; Vogel et al., 2013), but it is unknown whether myosin motors dwell at actin filament plus ends. There are intriguing observations of polar aster formation indicative of polarity sorting in cells (Verkhovsky et al., 1997) and in reconstituted assays (Backouche et al., 2006; Köster et al., 2016; Soares e Silva et al., 2011), but there is to our knowledge no direct evidence of myosin-mediated polarity sorting. Studies of contraction in actin-myosin networks have typically been performed at high protein densities, precluding direct observation of the interactions between actin and myosin at the single filament level.

Here we study how interactions between actin and myosin-2 at the single filament level translate into contractile activity at the network scale by performing time-lapse fluorescence imaging on reconstituted quasi 2D-networks of actin filaments and myosin motors. This 2D geometry mimics the quasi-2D random organization of the actin cortex and furthermore facilitates high resolution imaging of actin-myosin interactions at the single-filament level. Since actin-myosin remodeling is rather fast, we develop an open chamber assay that allows us to capture the initial steps of myosin-mediated remodeling immediately following the addition of components that trigger contractile activity. We show that myosin-2 bipolar filaments remodel initially random actin meshworks into polar asters. Experiments at low filament densities reveal that single myosin filaments processively walk towards actin filament plus ends, where they dwell. This end-dwelling behavior allows myosin filaments to transport actin filament ends together to form asters. By observing actomyosin remodeling over a range of time and length scales, we show that the polarity sorting at the single filament level drives aster formation and subsequent local or global network contraction, depending on the network connectivity. We use computer simulations to estimate the importance that polarity sorting and buckling may have for contraction *in vivo*, where the filaments are shorter and the network is denser than in our experiments *in vitro*.

## Results

To resolve the mechanism by which myosin drives remodeling of actin networks, we reconstitute two-dimensional networks of skeletal muscle actin and myosin labeled in different fluorescent colors on nonadherent glass surfaces and observe remodeling by total internal reflection fluorescence (TIRF) microscopy. Myosin-driven remodeling is a fast process whereby initial changes to the actin network happen on second time scales. In traditional flow cell setups these events would be difficult to capture. Therefore we use an open chamber setup, which allows us to trigger remodeling by adding myosin or ATP during the time-lapse acquisition (Fig. 1a, Video 1). When we add myosin to a pre-polymerized, random actin network, we observe that myosin starts to remodel actin as soon as it reaches the imaging surface (Fig. 1b). Within 40 seconds, this remodeling leads to the formation of actin asters with dense myosin foci at their center. Time-lapse imaging of the trajectories of myosins shows that they move directionally inward toward the center of the aster, implying that the asters are polar with actin plus ends oriented inwards (Fig. 1c and 1d). We conclude that myosin can remodel an initially random network of actin filaments within seconds into an organized network containing polar structures.

**Fig. 1.**
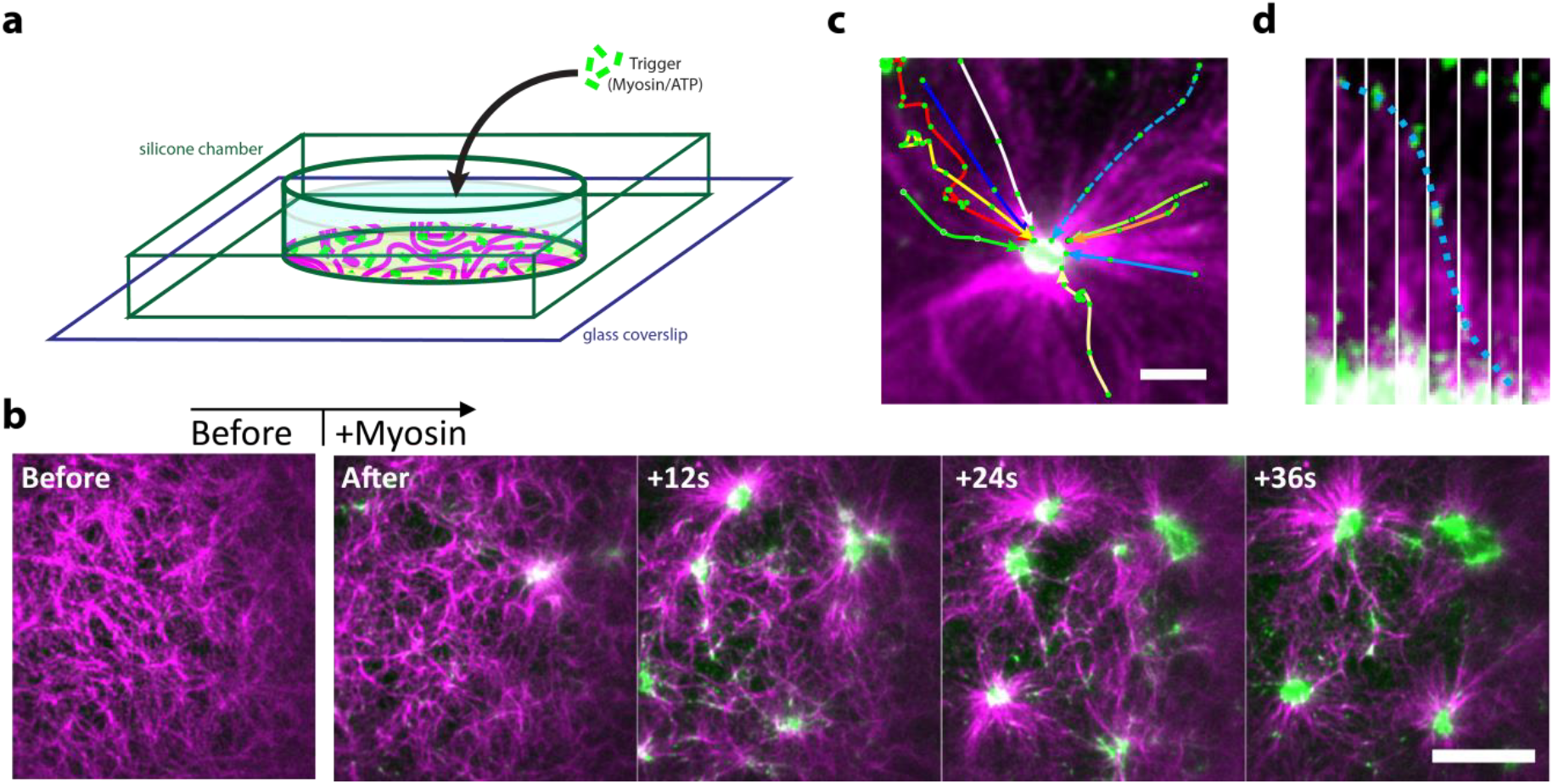
Myosin-induced actin network remodeling triggered by myosin addition. (a) An open chamber allows myosin or ATP to be added while the network located near the glass bottom is observed by time-lapse TIRF. (b) The initially disordered actin network (magenta) is rapidly remodeled by myosin (green) into asters. Scale bar is 20 μm and seconds after myosin addition are noted in the upper-left corner of the images. (c) Still image of an aster superposed with myosin trajectories (green data points connected by lines) measured over 206 s. Myosins move to the center of the aster indicating that actin filaments are polarity sorted with their plus ends oriented inwards. Scale bar is 5 μm. (d) Kymograph showing a myosin filament moving to the center of the aster. The trajectory corresponds to the blue dashed arrow in panel (c). Time between frames is 2 s.

To understand how the motors achieve polarity sorting, we used more diluted conditions where we can observe interactions of single myosin bipolar filaments with single actin filaments. We trigger myosin activity by adding ATP to a network of pre-polymerized actin and myosin. Prior to ATP addition, the actin filaments are densely decorated with myosin (left-most panel in Fig. 2a, Video 2), which is known to bind strongly in the absence of ATP. Under this condition, the actin filaments form bundles, likely due to the combination of the myosins acting as crosslinkers and the presence of a crowding agent. Upon ATP addition, most myosin filaments release, consistent with the low duty ratio of myosin-2 (Harris and Warshaw, 1993). The actin bundle disassembles in single filaments, likely due to the absence of crosslinking myosin. The remaining myosin filaments unidirectionally run along the actin filaments with a typical mean speed of 2 μm/s (Fig. 2b). We do not observe myosin detachment from the actin filaments unless the myosin filament encounters another actin filament, whereafter it switches track and runs along the new actin filament (after 28 seconds in Fig. 2a). As myosin reaches the end of the actin filament, it dwells on it. This can lead to an accumulation of myosin if several myosins run along the same actin filament (Fig. 2c, Video 3).

**Fig. 2.**
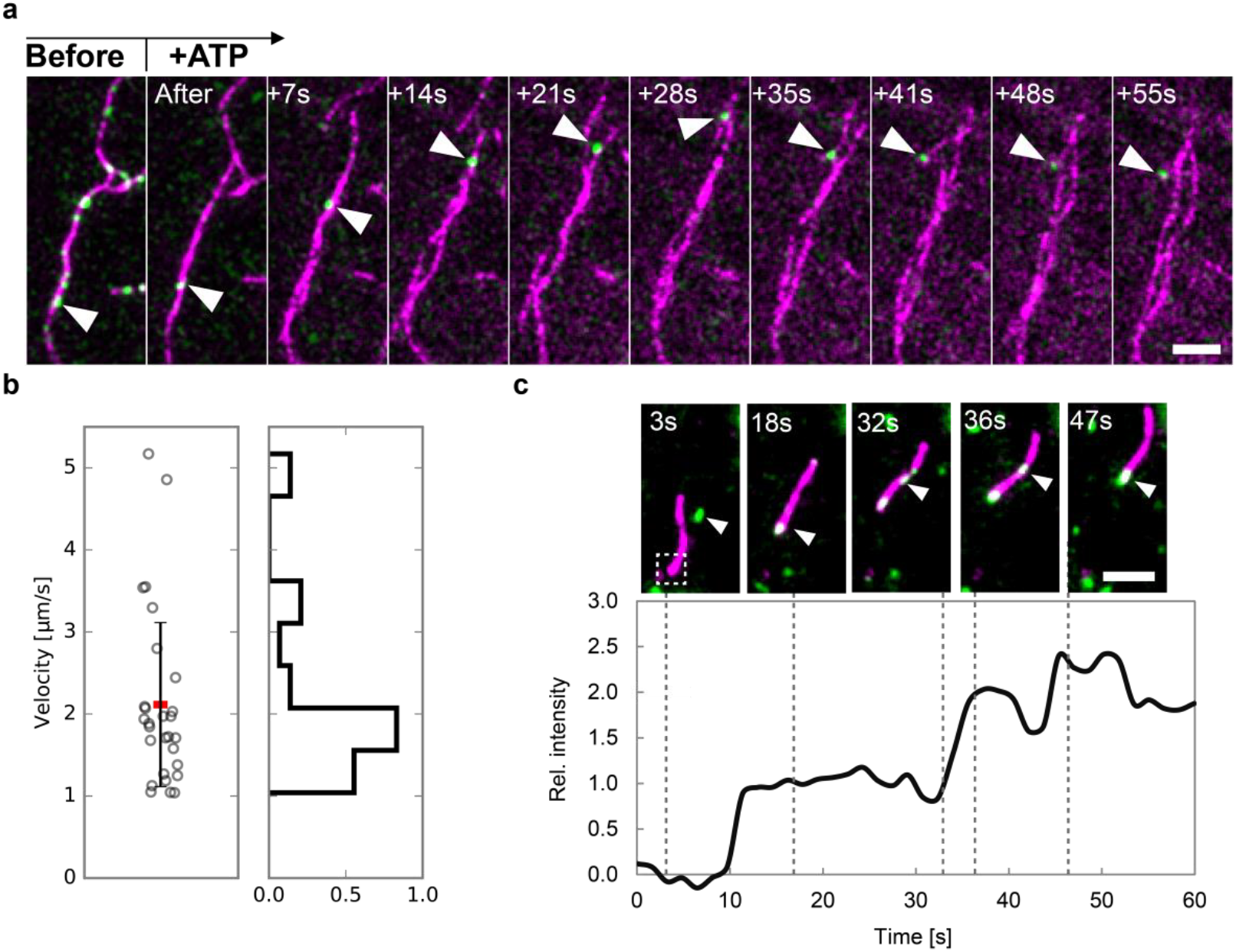
Myosin-actin interaction at low filament concentration. (a) Before ATP is added (left panel), myosin binds to actin and is immobile. Upon addition of ATP, most myosin filaments detach; one remaining filament runs along an actin filament, switches track as it encounters another actin filament at 28s, and runs on the new filament until it reaches the end and dwells there. Scale bar is 5 μm. (b) Left: velocities of myosins running along actin. Every data point corresponds to one myosin trajectory (n = 28). The horizontal red bar indicates the mean value (2 μm/s) and the error bar the standard deviation (1 μm/s). Right: Distribution of velocities. The velocity was only measured while myosin is moving.(b) Myosin accumulation at actin filament ends. Top: Three myosin filaments bind to an actin filament and move to the end. Bottom: Increase of fluorescence (Alexa Fluor 488) at the end of the actin filament (measured in the region indicated by white box in first panel above). The intensity is normalized to the intensity of the first myosin filament that reaches the actin end. Scale bar is 5 μm.

Myosin end-dwelling on single actin filament ends is a common event (Fig. 3a). This configuration is stable for at least several minutes. To determine a lower limit for the end dwelling time, we observed myosin end-dwelling for at least 4 minutes. In only 6% of the cases (n = 36), myosin detached during this time. From this we estimate a characteristic dwell-time of 64 min, assuming a Poisson process (see SI). We also find end-dwelling events that last over 15 min. Interestingly, myosin and actin ends seem not to overlap completely, which becomes clearer when we plot line profiles of the myosin and actin intensities (Fig. 3b). By analyzing these line profiles, we infer the length of the myosin filaments and overlap of the myosin and actin filament. We determine the average myosin length as 0.8 μm (Fig. 3c), in agreement with transmission electron microscopy data (Suppl. Fig. 1). The average overlap of the myosin and actin filaments is only 50% (Fig. 3c). This suggests that myosin filaments exhibit end-dwelling because of their “bipolarity” as sketched in Fig. 3d, consistent with prior reports of an interaction of F-actin with the trailing end of the bipolar myosin filament (Sellers and Kachar, 1990; Yamada and Wakabayashi, 1993).

**Fig. 3.**
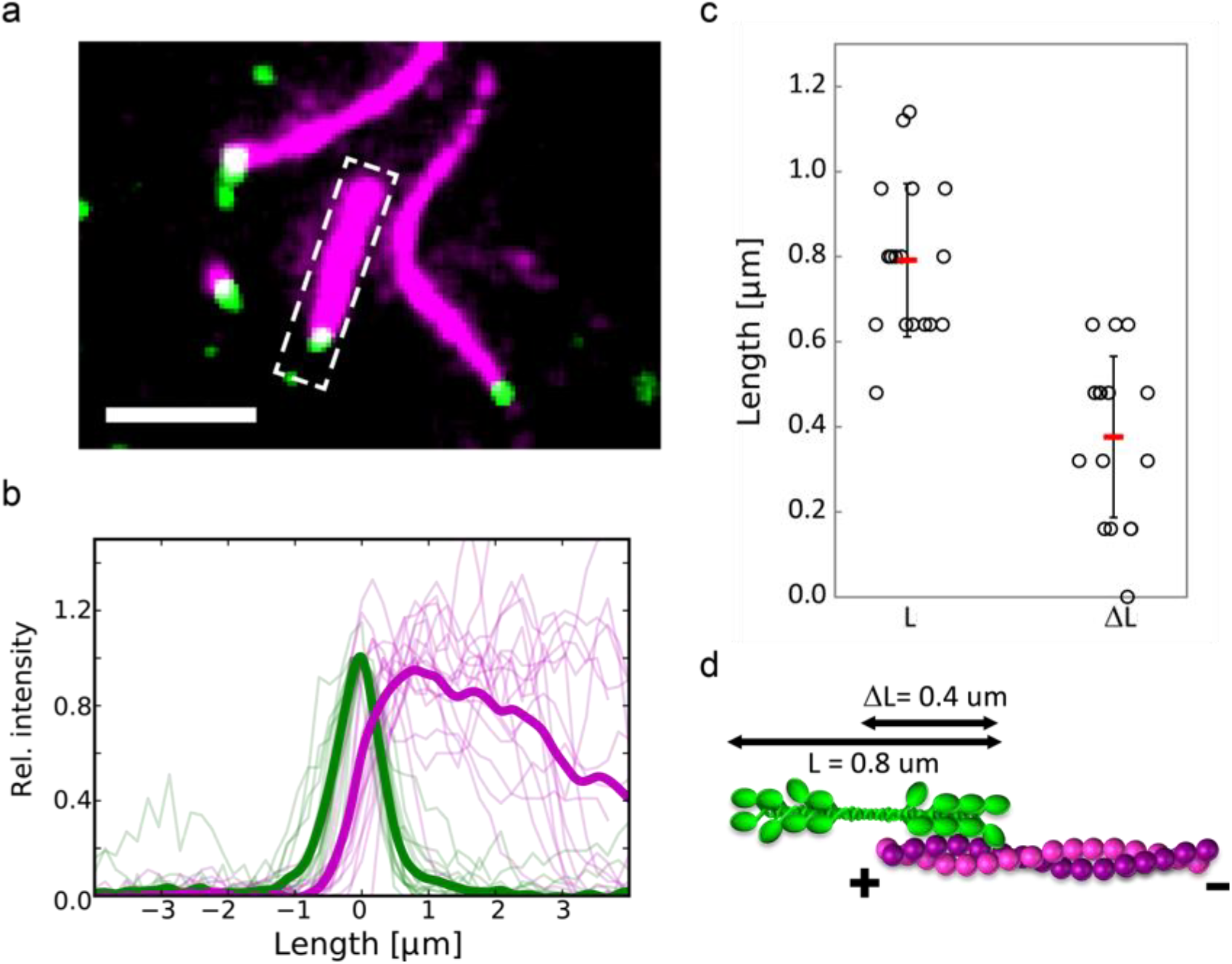
Position of myosin at actin filament ends. (a) Examples of myosin (green) dwelling on actin (magenta) filament ends. Scale bar is 5 μm. (b) Intensity line profiles of actin and myosin for 17 events (thin lines). Pairs of actin and myosin profiles are aligned with respect to the myosin peak intensity. The intensities are normalized to the peak intensity for each profile. Thick lines show the average profiles. (c) From the line profiles, we extract an average myosin filament length of 0.8 μm ± 0.2 μm (left) and an average overlap between end-dwelling myosin and actin filaments of 0.4 μm ± 0.2 μm (right); red bars indicate the mean values, error bars the standard deviation. (d) Schematic representation of myosin end-dwelling. As myosin only overlaps by about half of its total length, we suggest that the trailing end is involved in end-dwelling.

We then wondered whether the processive motion and end-dwelling behavior of myosin can account for the formation of actin asters via polarity sorting, given that theoretical models and simulations predict that these are necessary and sufficient ingredients (Surrey et al., 2001). Therefore we increased the actin filament concentration so that filaments overlap. We again observe that end-dwelling is a common event (Fig. 4a, yellow circles). Furthermore we see many examples of myosin connecting filament ends to form incipient asters (Fig. 4a, cyan circles). To understand the formation of these structures, we turned to live imaging. We see that myosin that is already bound to the end of one actin filament can still processively run along another actin filament (Fig. 4b, Video 4). As it arrives at the other filament’s end, it dwells also there and thereby holds the filament ends together. These bipolar structures can grow further and catch additional actin filaments (Suppl. Fig. 2). This observation shows that myosin end dwelling together with its processivity can be a mechanism to transport filament ends toward each other, as sketched in Fig. 4c. At concentrations of filaments and myosin used in this assay, the incipient aster configurations are not stable. Myosin can detach from one of the filaments while staying attached to the other (Fig. 4b, last time point). We used higher actin filament concentrations to test whether this mechanism is also active in dense networks (Fig. 4d, Video 5). We see the same mechanism: As myosin runs along one filament (cyan), it encounters a second filament (yellow). Subsequently it moves on both filaments and stops when it arrives at the two filament ends. This configuration is stable for a few minutes until myosin switches to another filament and starts to move on that one (Fig. 4d, last time point). Taking all this together, we conclude that myosin end dwelling together with processive motion can transport filament ends together, connect them in transiently stable complexes, and thereby induce polarity sorting.

**Fig. 4.**
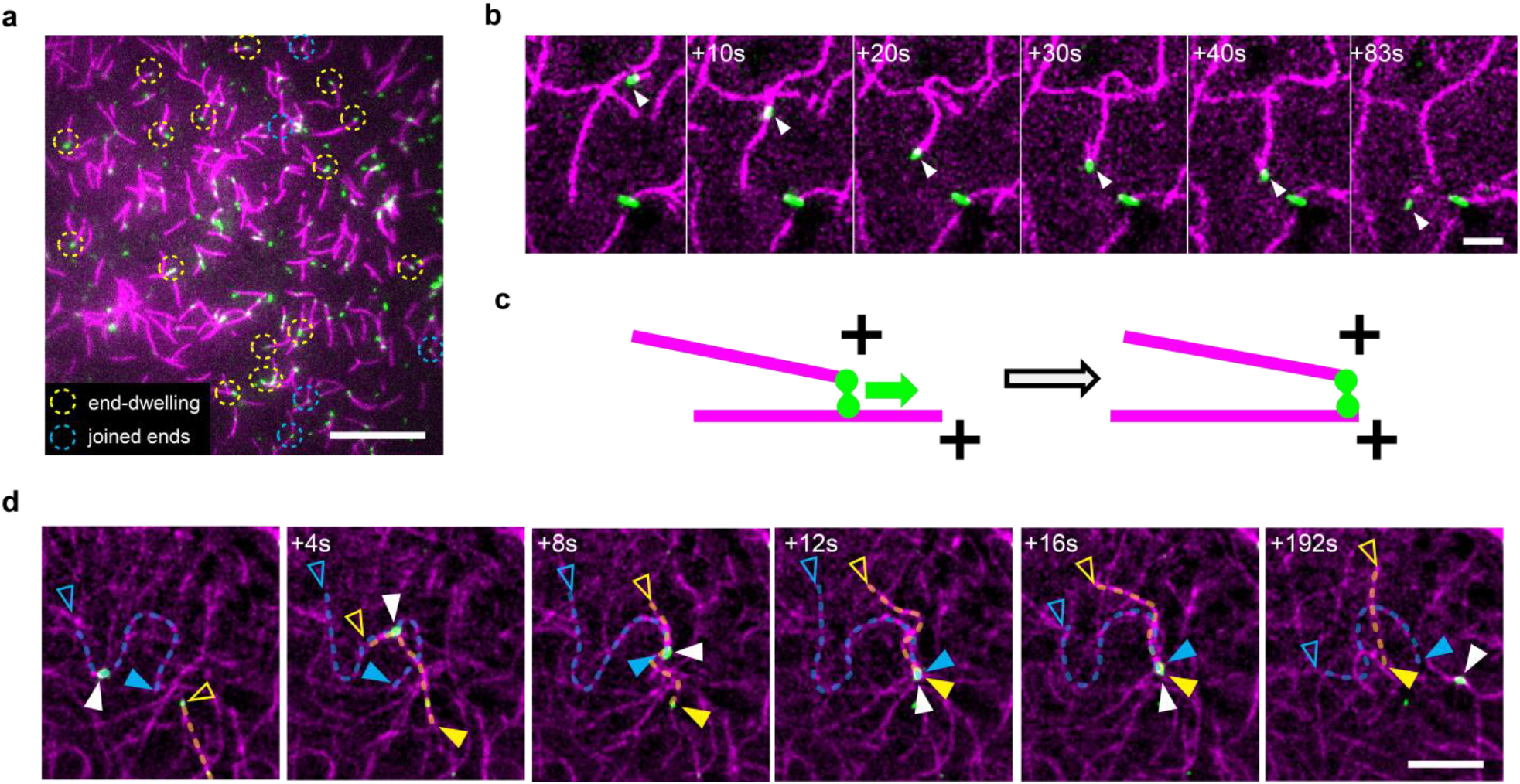
Myosin motility and end-dwelling as a mechanism for polarity sorting. (a) Myosin dwelling on actin ends (yellow circles) or connecting two filaments by their ends (cyan circles) in in conditions favoring sparse actin networks (see Table S1 for conditions). Scale bar is 20 μm. (b) Incipient asters forming in networks with intermediate numbers of filaments actin filament. Myosin attached to a short actin filament at t=0 encounters another actin filament at 10s and runs along it while staying attached to the first one, keeping the two filament ends together until it detaches from one at 83 s. Scale bar is 5 μm. (c) Schematic representation of the process of myosin-driven polarity sorting. Dwelling on one filament end, myosin transports the plus end as it runs along another filament and eventually joins the two plus ends. (d) Aster formation in networks with higher numbers of filaments. Myosin (white arrow head) runs along a filament (blue arrowheads and dashed line), encounters a second filament (yellow arrow heads and dashed line), and moves along both filaments, joining their plus ends. This configuration is stable for several tens of seconds. Scale bar is 10 μm, filled/empty arrow heads indicate plus/minus ends.

We next tested whether polar structures are also present at actin densities studied in prior reconstitution studies of contractility in actin-myosin networks (Murrell and Gardel, 2014, 2012; Vogel et al., 2013). We observed that asters still form and are rather stable (Fig. 5a), although we occasionally observe events where the aster splits (Fig. 5c, Video 6). The splitting of asters could be linked to the ability of the myosins to detach from filament ends when they get the option to switch to another actin filament track, as we observed at low filament densities.

**Fig. 5.**
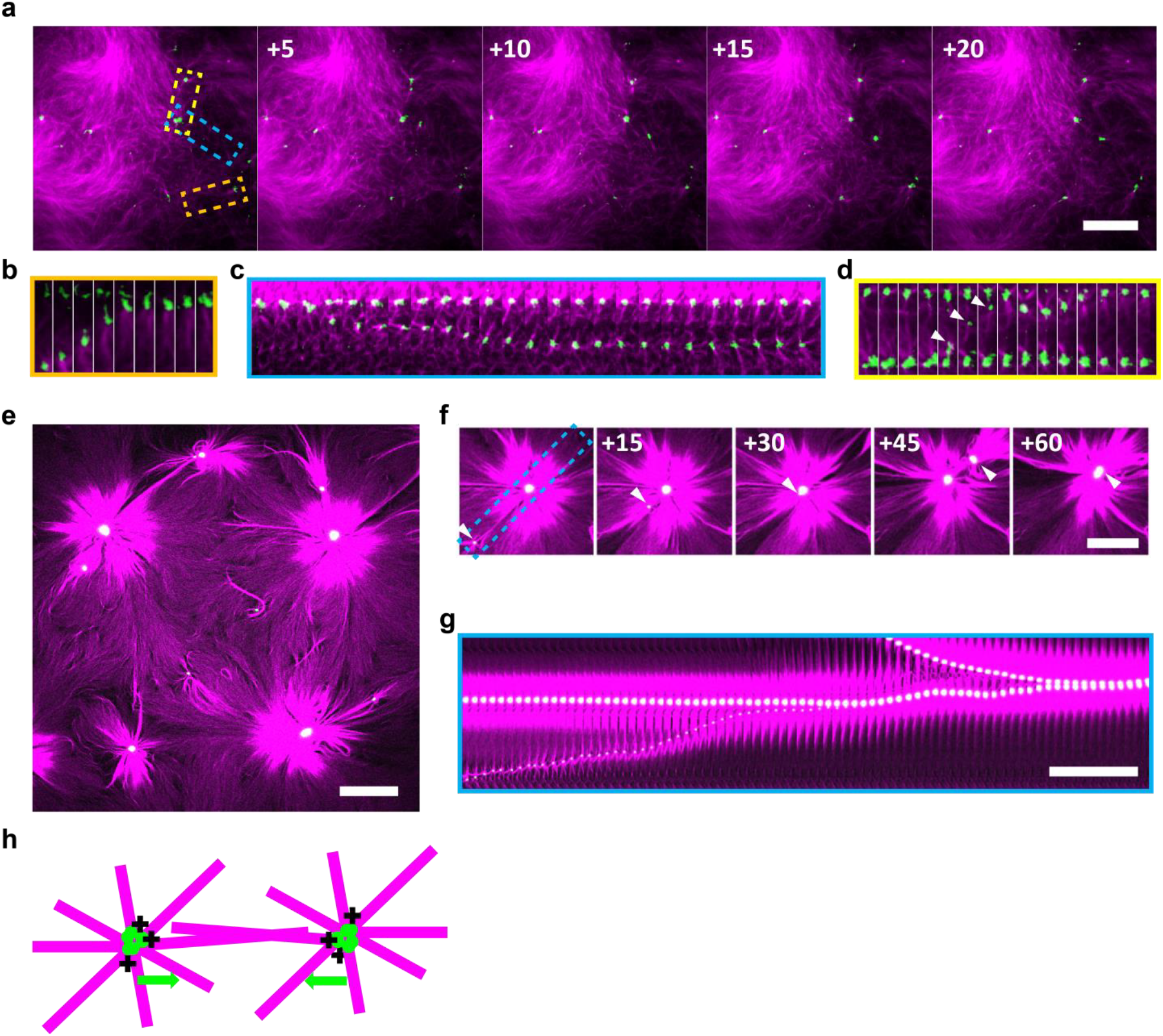
Effect of α-actinin-1 on myosin-mediated actin network remodeling. (a) In the absence of α-actinin-1, myosin (green) forms small actin asters (magenta). Time in seconds, scale bar is 20 μm. (b, c, d) Kymographs corresponding to the regions indicated by rectangles in panel (a) in corresponding colors. Time between frames = 1 s. (b) Joining of two asters. (c) Aster splitting in two. (d) Exchange of myosin between asters (arrow head). (e) In the presence of α-actinin (RC = 0.01), myosin remodels actin into large asters. Scale bar is 50 μm. (f) Two subsequent events where neighboring asters move towards each other in a persistent manner and eventually join. Time in min, scale bar is 50 μm. (g) Kymograph along the dashed line in (f). Scale bar is 10 min. (h) Schematic showing how neighboring asters move towards each other due to their polar nature.

Since asters are polar, they should in principle be able to merge due to the action of myosin motors. We indeed observe that neighboring asters interact. An example is shown in Fig. 5b: The two asters move toward one another but also away from each other along the same line. This one dimensional back and forth movement is clearly visible in the kymograph representation. Eventually the asters in this example merge. Interestingly we also observe exchange of myosin between asters (Fig. 5d).

We hypothesized that in order to increase aster interaction, we need to increase the network connectivity by adding a crosslinking agent. This hypothesis is based on evidence from *in vitro* studies, cells, and theoretical models that crosslinkers increase the range of force transmission by creating a percolated network (Alvarado et al., 2013; Bendix et al., 2008; Ennomani et al., 2016; Koenderink et al., 2009; Köhler et al., 2011; Laporte et al., 2012; Ojkic et al., 2011). Indeed, when we add the crosslinking protein α-actinin-1 (Ciobanasu et al., 2014), we observe that the asters are much larger, persistently move towards each other over distances of tens of μm, and invariably merge (Fig. 5e-g, Video 7). Long-range force transmission is mediated by α-actinin-rich actin bundles, which connect the asters (Suppl. Fig. 3). In addition to growing by merging, asters can also grow by a radial inward flow of actin filaments and myosin along the aster arms (Suppl. Fig. 4, Video 8). We conclude that crosslinkers favor contraction not only by providing elastic connections but also by allowing formation of stable F-actin bundle tracks that promote inward transport of myosin to form stable asters. Moreover, they form stable connections of mixed polarity between asters, stimulating aster merging (sketched in Fig. 5h). However, passive crosslinkers and motors acting together constitute the elementary configuration of the buckling mechanisms (Belmonte et al., 2017). Thus, in addition to helping the sorting mechanism, adding α- actinin is likely to promote the buckling mechanism as well.

We asked how this polarity sorting influences network contraction on large scales. As reported previously (Alvarado et al., 2013; Belmonte et al., 2017; Bendix et al., 2008), the balance between crosslinking and motor activity is a key parameter controlling the contractile behavior of actin-myosin networks. We fix the α-actinin concentration such that its molar ratio with actin is R_C_ = 1:50, which is well above the percolation threshold for 2D networks (Alvarado et al., 2017). To explore different regimes, we vary the myosin concentration. We confine actin-myosin networks containing 0.1 mM ATP in large ~2mm by 22mm chambers with non-adhesive walls and image the entire network over time using a low-NA (10x) objective, starting when contractility is triggered by mixing actin and myosin in the presence of ATP. Starting with a high myosin-to-actin ratio, R_M_ = 0.05, we find network contraction only on short length scales (Fig. 6a, magenta, Video 9). When we image the dense actin-myosin clusters at higher magnification, we find that they are comprised of actin asters with myosin foci at their core (Fig. 6b, magenta). As we decrease the motor-to-actin ratio R_M_ to 0.01, we still find actin asters but embedded in a fully percolated network as evidenced by a global network contraction (Fig. 6a). At higher magnification, we observe that the network is again made of asters (Fig. 6b). Decreasing R_M_ even further to 0.005 leads to stalled networks that are unable to contract (Fig. 6a, blue). Also here we observe asters at higher magnification (Fig. 6b). We suggest that two effects play a role in the contraction process. On the one hand, myosin accumulates in the center of the aster and is therefore depleted in the other areas, which do not experience major remodeling anymore. On the other hand aster arms have to overlap with neighboring aster centers to contract. These connections are more stable for lower myosin concentrations. As schematically depicted in Fig. 6c, our experimental results suggest that aster formation contributes to both local and global contraction.

**Fig. 6.**
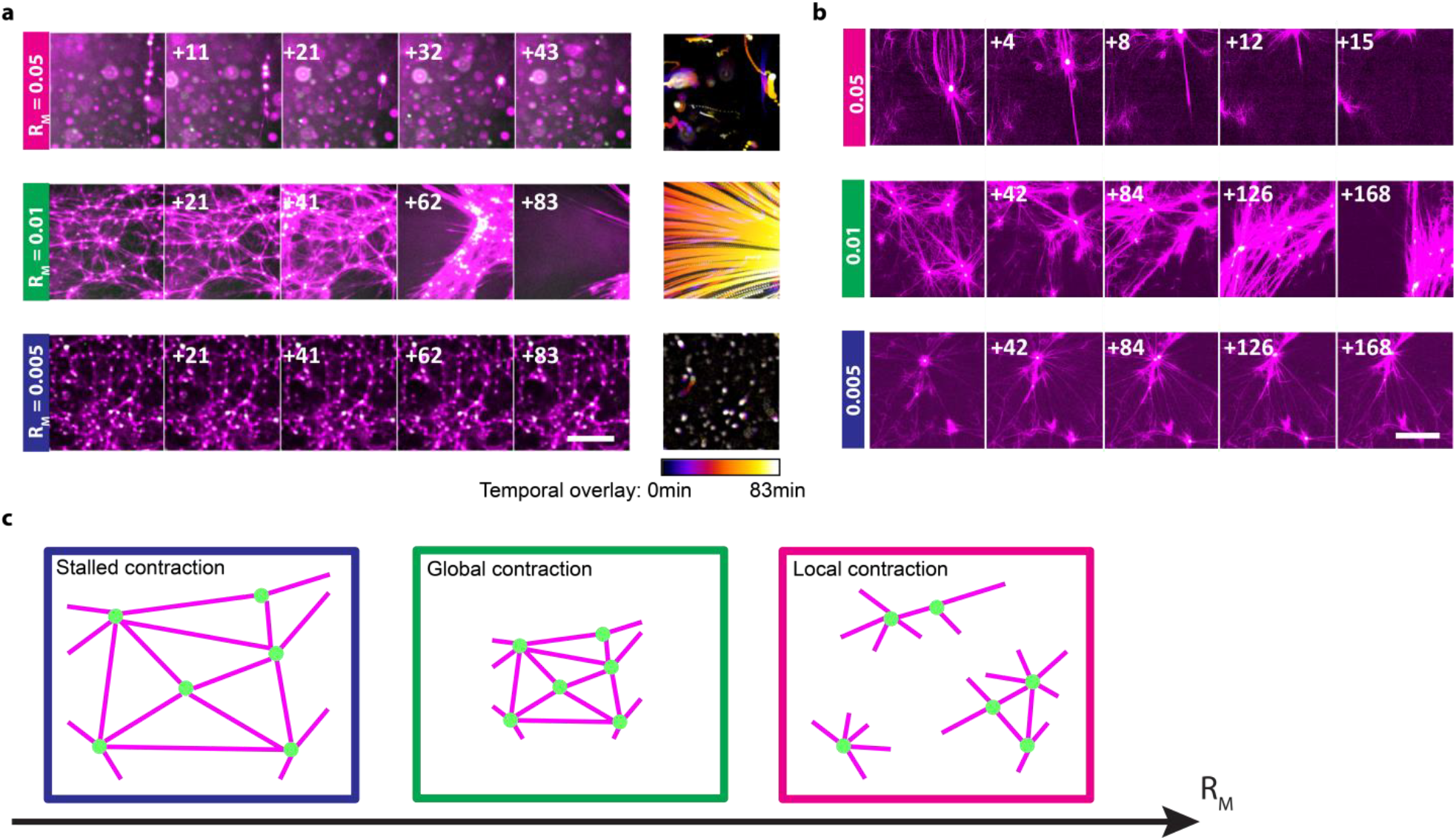
Asters formed at different motor:actin concentration ratios, RM. (a) At RM = 0.05, asters form and locally contract the network. For RM = 0.01, asters form and and become part of a percolating network that exhibits global contraction. For RM = 0.005, the network is stalled and does not contract, although asters are still present. Time in min, scale bar 200 μm. (b) Higher magnification images show that asters are formed in the three regimes. Time in min, scale bar 50 μm. (c) Schematic representation of the three contraction regimes as a function of RM.

While these large networks contract in the presence of both end-dwelling myosin-2 and α-actinin crosslinkers, it is not clear from the experiments if the observed contraction of the networks is due to a polarity-sorting mechanism alone or if the buckling mechanism or other effects are also working in parallel (Belmonte et al., 2017; Lenz et al., 2012; Ronceray et al., 2015). We can also not judge from our *in vitro* experiments to what extent this novel mode of contraction of actomyosin networks plays a role in physiological conditions, where both the filaments and mesh sizes are much smaller (Bovellan et al., 2014; Eghiaian et al., 2015; Fritzsche et al., 2016; Fujiwara et al., 2016). We thus turned to computer simulations of actin networks with the software Cytosim (Nédélec and Foethke, 2007), which allows us to approach *in vitro* and cortex-like conditions and analyze the relative contributions of end-dwelling and buckling mechanisms in networks of higher density.

We model individual filaments, myosin minifilament and α-actinin crosslinker explicitly, as described in the Methods section. While Cytosim has been used in the past to study both the buckling mechanism (Belmonte et al., 2017) and the polarity mechanism (Surrey et al., 2001) independently from each other, the current assumptions allow both mechanisms to operate in parallel here. Polarity sorting depends on the ability of the motors to end-dwell but does not require crosslinkers. Buckling-mediated contraction requires crosslinkers but operates even if the motors do not end-dwell. With simulations, we can thus enable or disable end-dwelling and add or remove crosslinkers to assess the influence of the two mechanisms. We first modeled contraction of actin networks with myosin minifilaments but no crosslinkers, with myosin dwelling at filament ends. The end-dwelling property of myosin heads leads to the formation of asters (Fig. 7a, Video 10), with all the plus ends facing inwards (Fig. 7a, last panel), resembling the asters observed *in vitro* (Fig. 1). If we let the motor heads detach immediately upon reaching the end of actin filaments, the network fails to contract (Fig. 7b, Video 11). These results confirm the role of end-dwelling for the contractility observed in the *in vitro* networks.

**Fig. 7.**
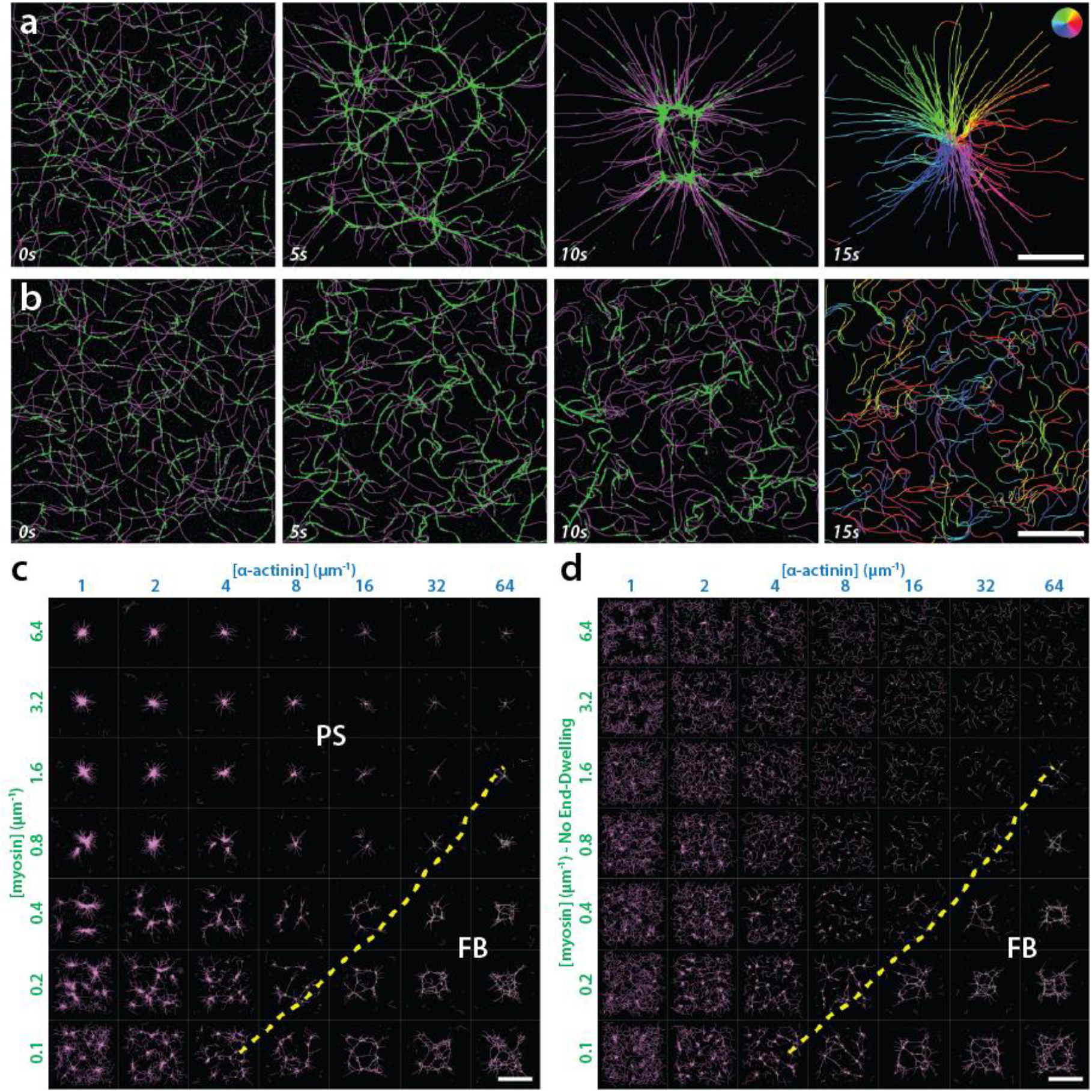
Role of end-dwelling in simulated networks with parameters corresponding to *in vitro* experimental conditions. (a, b) Time series of simulations with 200 flexible filaments (“actin”, in magenta) with average lengths of 10 μm, and myosin minifilaments (“myosin”, in green). Myosin motors are plus-end directed and modeled (a) with or (b) without end-dwelling. The last time point shows actin filaments segments colored according to their orientation. In (a) asters form with all filament plus ends at the center of the aster as a result of the polarity sorting mechanism. Without end-dwelling (b), the network does not contract. (a, b) Scale bar 10 μm. (c, d) Phase diagrams of simulated networks after 200s of simulation with varying concentrations of α-actinin and myosins (c) with or (d) without end-dwelling motors. Concentrations of α-actinin and myosin are given as molecules per μm of actin filament. Actin filament lengths as in (a, b) with mesh-size of 1.1 μm. In the region above the dashed yellow line, contraction is dominated by Polarity Sorting (PS), as shown by the fact that contraction is lost without myosing end-dwelling. In the region below, contraction persists also without myosin end-dwelling, showing it is mediated by Filament Buckling (FB). (c, d) Scale bar 40 μm.

We next added α-actinin crosslinkers and systematically varied the concentration of myosin and α–actinin (Fig. 7c). We can observe all three network behaviors seen in the *in vitro* experiments: slow or stalled networks, for high concentrations of α-actinin and low concentrations of myosin; globally contracted networks for high myosin concentrations; and locally contracted asters for low myosin and α- actinin concentrations. To assess the contribution of polarity-sorting to the contraction of the networks, we repeated the simulations with the end-dwelling turned off (Fig. 7d). In this scenario most of the networks fail to contract, indicating an important role of polarity sorting in the contraction of actin networks. The region of the phase diagram least affected by the absence of end-dwelling is characterized by high concentrations of crosslinkers, a condition that is crucial in the filament buckling mechanism, as shown previously by theory and simulations (Belmonte et al., 2017; Lenz et al., 2012) and experimentally (Bendix et al., 2008). For buckling to happen, the crosslinking density must be high enough to sustain the forces exerted by the myosin motors while buckling the filaments. Therefore, we expect the boundary between the filament buckling or the polarity sorting mechanisms to be determined by the ratio of end-dwelling myosins to crosslinkers. On the diagram where the two concentrations are varied, the regions dominated by either mechanism are thus separated along a diagonal (Fig. 7c, d).

The above results highlight the prominent role of polarity-sorting in the contraction of networks that resemble the *in vitro* conditions reported in our experiments. In physiological conditions, for example in the cortex of animal cells, the actin filaments are about one order of magnitude shorter than in our *in vitro* systems (about 1 μm or less (Fritzsche et al., 2016)), and the mesh size is also much shorter (between 0.03 to 0.1 μm (Bovellan et al., 2014; Fujiwara et al., 2016)). One can thus expect polarity-sorting to be even more critical in the cortex for two reasons: Firstly, the force required to buckle a filament segment increases with the inverse of the squared length of the segment, and thus becomes greater as the mesh size is reduced. Secondly, the ratio of end-dwelling versus side-bound motors would also increase as the filaments become shorter. To assess these effects, we performed simulations with denser networks made of shorter filaments that better resemble the actomyosin cortex and varied the concentrations of myosin and α-actinin as before (Fig. 8a). With these conditions, most of the networks contract to a single aster rather than multiple ones, probably because they are more tightly connected. When the end-dwelling property of myosin is turned off, the contractile behavior is lost in a region of the phase diagram that is similar to the region that lost contractions under the simulated conditions of the *in vitro* experiments (Fig. 7c, d). However, a significant number of networks with low myosin and high α-actinin concentrations are now stalled, whereas they were contractile under the buckling mechanism at lower density. Therefore, under conditions mimicking the actin cortex *in vivo*, the effectiveness of the buckling mechanism is reduced, while polarity sorting is preserved. Note that for low α-actinin concentrations, filaments freely glide over the network, powered by the myosin motors. In these conditions, filaments are pushed out of the initial geometry more readily, due to the smaller overall available space, and the effect is more visible compared to the *in vitro*-like simulations (Fig. 7d). These results confirm that the polarity sorting mechanism remains effective when the characteristics of the actomyosin networks are more cortex-like, while at the same time the buckling-mediated mechanisms decreases in importance.

**Fig. 8.**
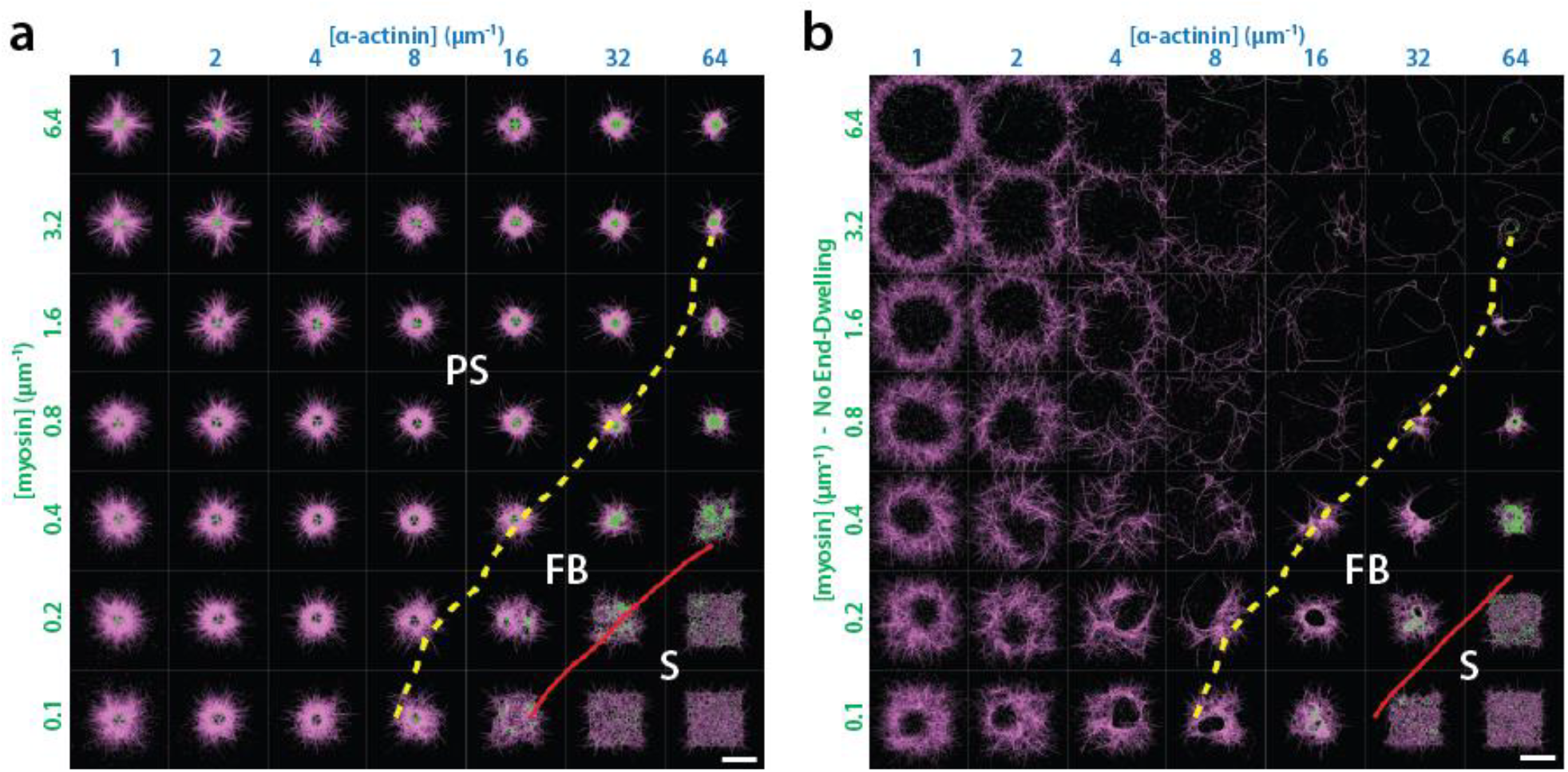
Role of end-dwelling in simulated networks with parameters corresponding to the actin cortex *in vivo*. (a, b) Phase diagrams of simulated networks after 80s of simulation with average actin filaments lengths of 1 μm, mesh-size of 0.1 μm and varying concentrations of α-actinin and myosins (a) with and (b) without end-dwelling motors. Concentrations of α-actinin and myosin are given as molecules per μm of actin filament. Dashed yellow lines demarcate the regions where contraction is dominated by Polarity Sorting (PS) or Filament Buckling (FB), as described in Fig. 7. Solid red lines indicate the boundaries between contractile and Stalled (S) regions in each phase diagram. Scale bar 4 μm.

## Discussion

An important biological function of actin-myosin networks is their contractility, for which the underlying mechanism is still not fully understood. Here we show that myosin motors remodel initially random actin networks into polarized domains. Using low protein densities to observe actin-myosin interactions at the single filament level, we identify myosin dwelling at actin filament ends as the mechanism for this polarity sorting. Our study thus reveals how contractile nodes, which are often assumed in theoretical studies of actin-myosin networks, form by self-organization (Alvarado et al., 2013; Hannezo et al., 2015; Jülicher et al., 2007; Salbreux et al., 2009). We find that aster formation occurs over a wide range of crosslinker and myosin densities. At low crosslink densities that are below the percolation threshold, the asters are relatively small and transient. At crosslink densities above the percolation threshold, the asters are stabilized and neighboring asters pull on each other over long distances and merge, leading to large-scale contraction. The crucial role for crosslinkers to tune the range of force transmission is in line with prior experimental and theoretical work (Alvarado et al., 2013; Bendix et al., 2008; Chugh et al., 2017; Vavylonis et al., 2008). But we find that crosslinkers favor contraction not only by providing elastic connections but also by allowing formation of stable F-actin bundle tracks that promote inward transport of myosin to form large and stable asters. By contrast, in the absence of crosslinkers, myosin clusters are dynamic and split because myosin filaments can switch between different actin filament tracks. Asters are also observed over a range of motor densities, in conditions of stalled networks (low motor density), globally contracting networks (intermediate motor density), and local contraction (high motor density). These contractile behaviors are reproduced in simulations, where we can directly compare the remodeling effect of motors with and without end-dwelling. The simulations demonstrate that end-dwelling is an important requirement for network contraction while contraction due to filament buckling only happens for a limited set of parameters

In our study we use skeletal muscle myosin-2 filaments. While muscle myosin is used in nearly all reconstituted actomyosin systems thus far, with few exceptions (Thoresen et al., 2013), contraction in other cells is driven by non-muscle myosins. These differ from skeletal muscle myosin 2 in their kinetic properties and the formation of much smaller bipolar filament ensembles of only 10-20 motors (Erdmann et al., 2016). Clearly, in future it will be interesting to study if end-dwelling is also observed for non-muscle myosin. Our results suggest that the dwelling behavior is mainly mediated by the trailing end of the myosin filaments. While non-muscle myosin filaments are reported to be much shorter than skeletal muscle myosin filaments (Billington et al., 2013), their longer duty ratio might compensate and still permit end-dwelling (Kovács et al., 2003; Wang et al., 2003). Possibly the ratio between the two major non-muscle myosin isoforms, NMMII A and B, could also play a role in the stability of filament end-dwelling (Melli et al., 2017).

We show that myosin-driven polarity sorting can be an efficient mechanism for actomyosin network contraction provided that enough crosslinkers are present to allow force transmission. Prior studies of reconstituted actin-myosin networks showed that in the presence of crosslinkers, motor-mediated buckling of actin filaments can provide the dominant mechanism for contraction, sometimes augmented by severing of buckled actin filaments (Murrell and Gardel, 2012). But both our experimental and computational findings suggest that even in crosslinked networks, there is still an underlying tendency for actin and myosin to form polarity-sorted structures. The relative contributions of polarity sorting versus buckling to the overall network contraction depend on microscopic parameters such as the distance between crosslinks, the motor density, and whether actin filaments are bundled. A full characterization of these parameters is beyond the scope of this study. Simple scaling considerations indicate that buckling may be disfavored in the *in vivo* actin cortex. First, recent work suggests that the actin cortex is composed of filaments that are relatively short, being a mixture of formin-nucleated filaments with lengths on the order of 1 μm, and Arp2/3-nucleated filaments in the 100 nm range (Fritzsche et al., 2016). Second, electron microscopy images revealed typical cortical mesh sizes of only 30 to 200 nm, which might correspond to the distance between crosslinkers (Bovellan et al., 2014; Fujiwara et al., 2016). These characteristics would make buckling difficult, but leave polarity sorting as a potent contraction mechanism, as our *in silico* study showed.

There are some reports of actin polarity sorting in a few cellular structures by electron microscopy (Begg et al., 1978; Cramer et al., 1997; Kamasaki et al., 2007; Sanger and Sanger, 1980). Furthermore, myosin foci are commonly observed in the actin cortex of cells and early embryos. However, the cell cortex is mainly regarded as a random actin network. Recent theoretical studies of contractile networks with actin turnover suggest that the absence of asters in the cell cortex may be due to turnover (Guthardt Torres et al., 2010; McFadden et al., 2017). Consistent with this idea, asters can be formed in the actin cortex if actin polymerization is reduced or blocked by drugs (Luo et al., 2013; Verkhovsky et al., 1997), and actin stabilization seems to have a similar effect (Wehland et al., 1977). We therefore speculate that polarity sorting may occur in cortical actin networks but that actin turnover prevents formation of large polar domains. Moreover, polarity sorting may be counteracted by other remodeling processes such as Arp2/3 nucleator-dependent self-organization (Fritzsche et al., 2017) and biochemical feedback between active RhoA and myosin (Nishikawa et al., 2017). New developments in high resolution microscopy will hopefully enable the visualization of local accumulation of actin plus ends in the cortex or other contractile structures such as the cytokinetic ring, and thereby lead to a better understanding of the role of polarity sorting in cells (Hu et al., 2017).

## Materials and Methods

### Protein preparation

Actin and myosin are purified from rabbit muscle as described previously (Alvarado and Koenderink, 2015). Myosin is labelled with Alexa Fluor 488 NHS ester (Invitrogen). Actin is labelled with Alexa Fluor 649 carboxylic acid, succinimidyl ester. The crosslinker protein α-actinin-1 in unlabeled form and tagged with mCherry is purified as described in (Ciobanasu et al., 2014). Myosin is stored in myosin buffer (300 mM KCl, 4 mM MgCl_2_, 20 mM imidazole, 1 mM dithiothreitol (DTT)) with 50% glycerol. Prior to the experiment, myosin is dialyzed overnight against the myosin buffer to remove the glycerol. At this salt concentration myosin does not self-assemble into filaments. Actin is stored in G-Buffer (2 mM Tris-HCl, 0.2mM ATP, 0.2 mM CaCl_2_, 0.2 mM DTT) in its monomeric (G-actin) form.

### Chamber preparation

Glass coverslips are cleaned with base piranha for 10 min(Alvarado and Koenderink, 2015) and thoroughly rinsed with milliQ water. A homemade silicone sheet with 9mm^2^ holes is placed on top of the glass coverslip to build open chambers. In some experiments (Table 1, supplementary information), classical flow cells are used similar to the ones described in (Alvarado and Koenderink, 2015). The surfaces are passivated with lipid bilayers: small unilamellar vesicles (SUVs, see below) are flushed into the chamber. After an incubation time of at least 5 min, excess vesicles are flushed out with F-buffer (50 mM KCl, 2 mM MgCl_2_, 20 mM imidazole). To prevent drying between flushing steps, the chamber is kept in a humid atmosphere.

### SUV preparation

Lipids are stored in chloroform. To remove chloroform, lipid solution (typically 50 μl) is pipetted into a glass tube. Chloroform is slowly evaporated by gentle nitrogen flow while turning the tilted glass tube to achieve a homogenous layer of lipids at the bottom of the tube. To remove any remaining chloroform, the tube is kept in vacuum overnight. The lipids are resuspended in buffer (20 mM imidazole, 50 mM KCl) and sonicated with a tip sonicator for 30 min to make small unilamellar vesicles (SUVs). We use 1.9 mg/ml of the neutral lipid DOPC (1,2-dioleoyl-sn-glycero-3-phosphocholine, Avanti Polar Lipids). For open chamber experiments we supplement the membrane with 1 mol% of PEGylated lipids PEG-PE (1,2- dipalmitoyl-sn-glycero-3- phosphoethanolamine-N-[methoxy(polyethylene glycol)-2000], Avanti Polar Lipids) to make the bilayer more resilient against drying.

### Contraction experiment

In the cases where we use an open chamber, actin is prepolymerized for longer than 1h. Myosin is then added with the contraction buffer (50 mM KCl, 2 mM MgCl_2_, 20 mM imidazole, 1 mM DTT, 0.1 mM ATP, 2 mM protocatechuic acid (PCA), 0.1 μM protocatechuase 3,4-dioxygenase (PCD), 10 mM creatine phosphate (CP), 0.1 mg/ml creatine kinase (CK), 0.3 % methyl cellulose (MC)). CP and CK are used as an ATP replenishing system, while PCA and PCD are used as an oxygen scavenger system and MC acts as a crowding agent that pushes actin and myosin filaments towards the coverslip. In flow cell experiments, monomeric actin is mixed with the contraction buffer, which triggers its polymerization, and the mix is immediately injected into the channel. Exact actin and myosin concentrations can be found in Table 1 of the supplementary information.

### Diluted filament assay

Diluted filament conditions shown in Fig. 2 to 4 are obtained either by polymerizing actin is at low concentrations or by slowing down polymerization of an initially dense actin solution by diluting after a few minutes. The exact conditions can be found in Table 1 of the supplementary information. Myosin is added to the filaments in the contraction buffer. For experiments where myosin activity is triggered by addition of ATP solution to a final concentration of 0.1 mM (Fig. 2a), the contraction buffer does not contain ATP.

### Microscopy and image analysis

Confocal images were taken with a Nikon Eclipse Ti inverted microscope equipped with a Nikon C1 confocal scan head and a 100-mW Argon ion laser (488 nm, 561 nm, Coherent, CA), or with a Nikon Eclipse Ti inverted microscope equipped with a CrEST spinning disk unit, a solid state light source (SpectraX, Lumencor, OR) and a Hamamatsu camera. Typical exposure time was 200 ms. Total internal reflection fluorescence (TIRF) imaging was performed with a Nikon Eclipse Ti-E inverted microscope equipped with a Roper TIRF module and QuantEM:512SC EMCCD camera. Exposure time was 100 ms.

Image analysis was performed with the ImageJ distribution Fiji (Schindelin et al., 2012; Schneider et al., 2012). The intensity background was subtracted with Fiji. For representation purposes images were smoothened. The velocity of myosin filament motion on actin filaments (Fig. 1c and 2b) was measured by manual tracking of single myosin filaments using the “Manual Tracking” plugin of ImageJ. Temporal overlays (Fig. 6, Suppl. Fig. 4) were made with the ImageJ plugin “Temporal-Color Coder”. Kymographs were made with Fiji. To detect myosin and actin filament edges (Fig. 3), we took line profiles of the respective fluorescence signals. The edge position was determined as the maximum in the first derivative of the profiles.

### Electron microscopy

Transmission electron microscopy images were taken with a FEI Verios 460. Myosin was assembled in F-Buffer at 0.2 μM. The solution was diluted (typically 1:100), applied on an electron microscopy grid (300 mesh Cu grid, Ted Pella inc., CA), and rinsed with ultrapure water after 1 min of incubation. Finally the grid was air dried. Myosin length measurements were performed manually with the ImageJ distribution Fiji (Schindelin et al., 2012; Schneider et al., 2012).

### Computer Simulations

The simulations of contractile actin-myosin networks were performed with the Open Source software Cytosim (github.com/nedelec/cytosim), which uses a Brownian dynamics approach as described previously (Nedelec and Foethke, 2007). Actin filaments are modeled as incompressible bendable filaments of rigidity 0.075 pN.μm^2^ (corresponding to a persistence length of 18 μm) in a medium of viscosity 0.18 Pa.s. The α-actinin crosslinkers are modeled as Hookean springs of zero resting length and a rigidity of 50 pN/μm, with a binding rate *k*_on_ = 15 s^-1^, a binding range of 0.02 μm and a slip-bond unbinding model: *k*_off_ = *k*_off,0_ exp(*-|f|*/*f*_0_) with a basal unbinding rate of *k*_off,0_ = 0.3 s^-1^ and unbinding force of *f*_0_ = 2 pN. Myosin mini-filaments were modeled as an inextensible 1D object of length 0.8 μm with 4 motors on each side spaced by 0.08 μm from the extremities. Motors operate independently from each other and also behave as a Hookean spring when attached to an actin filament, with zero resting length and a rigidity of 100 pN/μm. Additionally motors move on filaments with a speed of 2 μm/s, with a linear force-velocity relationship characterized by a stall force of 4 pN. Each motor has a binding rate of *k*_on_ = 10 s^-1^, a binding range of 0.01 μm and a force-independent unbinding rate of *k*_off_ = 0.5 s^-1^. Unless otherwise specified, motors end-dwell by stopping upon reaching the end of filaments without changing their unbinding rates. Motors and α-actinin binding and unbinding events are modeled as first-order stochastic processes and we neglected steric interaction between filaments, motors and crosslinkers.

The simulations without α-actinin (Fig. 7a-c) were done with 200 actin filaments randomly distributed over a square area of 40x40 μm^2^, with filament lengths following an exponential distribution with a mean of 10 μm that was truncated inside [0.5; 20] μm. For the simulations reproducing the *in vitro*-like conditions (Fig. 7d,e), 800 actin filaments were randomly distributed over a square area of 80x80 μm^2^, with filaments lengths distribution as before. In those simulations the resulting mesh-size (measured as the average distance between consecutive filament crossings) was about 1.1 μm. For the simulations that reproduce the cortex-like conditions, 3500 filaments were randomly distributed over a square area of 8x8 μm^2^, with exponentially distributed lengths, with a mean of 1 μm truncated in [0.1; 4] μm. The resulting mesh-size was about 0.1 μm. In both scenarios the filaments are mixed with varying amounts of α-actinin crosslinkers and myosin mini-filaments. The concentrations of myosin and α-actinin are defined as number of elements per total length of actin (units of μm^-1^) and were varied within the ranges [0.1:6.4 μm^-1^] and [1:64 μm^-1^], respectively, for both scenarios. To better mimic experimental conditions, actin filaments are not created as straight filaments, but already relaxed according to their persistence length (18 μm), and the simulations start with all motors and crosslinkers unbound. The reference configuration files for each scenarios are provided as Supplementary File.

In the phase diagrams (Figs. 7d,e and 8a,b), the boundary between the Polarity Sorting (PS) and Filament Buckling (FB) mechanisms was determined by comparing the contraction rates of the with and without end-dwelling. The contraction rates were calculated as in (Belmonte et al., 2017). The regions of the phase diagram where 50% or more of the contraction is lost when end-dwelling is turned off were considered to be dominated by the Polarity Sorting mechanism. The boundary line is estimated by creating a matrix with the normalized difference in contraction between the simulations with and without end-dwelling, which was later smoothed using a 2D boxcar average with window size of 3. We then used the matplotlib library from python to calculate the isocline at 0.5. The networks that experience a small initial contraction (10 % or less) that stopped after a few seconds were considered to be non-contractile and made the Stalled (S) region of the phase diagrams of Fig. 8. Note that the networks in same region of the phase diagrams in Fig. 7 experience a slow but continuous contraction, and therefore were not considered stalled.

## Acknowledgements

This work is part of the research program of the Netherlands Organisation for Scientific Research (NWO) and was financially supported by an ERC Starting Grant (335672-MINICELL). We thank Dr. C. Le Clainche (Institute for Integrative Biology of the Cell (I2BC), Université Paris-Sud) for kindly providing the α- actinin-1 constructs and Marjolein Vinkenoog-Kuit for protein purification. FJN is supported by the Center for Modelling and Simulation in the Biosciences (BioMS). JMB and ML are supported by EMBO and EMBL. We wish to acknowledge EMBL IT support.

